# Beyond the Motor Cortex: Theta Burst Stimulation of the Anterior Midcingulate Cortex

**DOI:** 10.1101/707802

**Authors:** Travis E. Baker, Mei-Heng Lin, Seema Parikh, Neeta Bauer, Carrisa Cocuzza

## Abstract

Intermittent Theta Burst Stimulation (TBS) applied to the left dorsolateral prefrontal cortex suppressed reward-related signaling in the anterior midcingulate cortex (aMCC), resulting in a change in goal-directed behavior. Continuous TBS had no effect. While these results are inconsistent with reported TBS effects on motor cortex, the present findings offer normative insights into the magnitude and time course of TBS-induced changes in aMCC excitability during goal-directed behavior.

---

Theta-burst stimulation (TBS) protocols have recently emerged as having a fast and robust faciliatory (intermittent TBS: iTBS) and inhibitory (continuous TBS: cTBS) action on motor cortex excitability^1^. As a result, several groups have begun exploiting its potential in the study of prefrontal function in both typical^2^ and atypical^3^ populations. While TBS effects on motor cortical excitability can be assessed relatively directly by measuring motor‐evoked potentials, the evaluation of TBS when applied to prefrontal regions requires different approaches, such as PET, EEG, and fMRI. However, combined TBS neuroimaging studies designed to investigate the after-effects of TBS beyond the motor cortex have largely produced conflicting results^4,5^. Collectively, the action of TBS protocols on neurocognitive functioning remain unclear. Here, we used robot-assisted TBS, in combination with scalp electrophysiological recordings, to examine whether TBS protocols applied to the left dorsal lateral prefrontal cortex (DLPFC) can differentially modulate the activity of the anterior midcingulate cortex (aMCC) during goal-directed behavior.

The function of aMCC is hotly debated, but an influential theory holds that aMCC utilizes reward prediction error signals (RPEs) for the purpose of reinforcing adaptive behaviors. In humans, this process is revealed by a component of the event-related brain potential called the reward positivity^6^ (Figure 1C). Converging evidence indicates that the reward positivity is generated in aMCC and indexes an RPE signal^6^. In parallel, 10-Hz repetitive transcranial magnetic stimulation (rTMS) applied to the left DLPFC has been shown to enhance dopamine release^7^, neuronal activity, and cerebral blood flow^8^ in the aMCC, as well as increase the amplitude of the reward positivity^9^. Leveraging these two streams of evidence, we first asked whether iTBS and cTBS applied to the left DLPFC could facilitate or disrupt aMCC activity, as evaluated by the reward positivity. If true, we then asked whether a TBS-induced change in aMCC activity would cause a change in goal-directed behaviour (e.g. win-stay and lose-shift actions). Given the importance of the aMCC to normal and pathological function, it would be a great theoretical and therapeutic advantage if one could specifically drive or inhibit aMCC activity with TBS.

**Figure 1.**
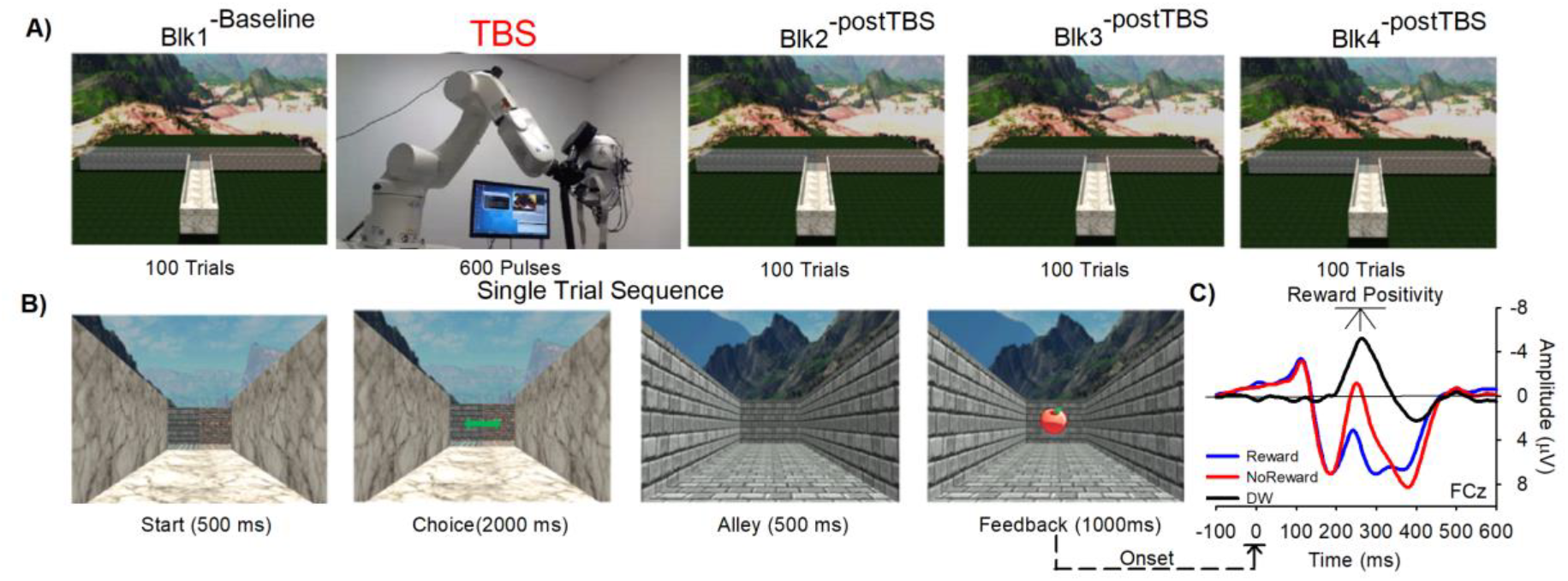
TBS paradigm. A) Block sequence. B) Single-Trial Sequence. C) ERPs elicited by reward feedback (blue), no-reward feedback (red) and difference wave (black). The reward positivity is observed as a differential response in the ERP to reward and no-reward feedback, occurring over frontal–central areas of the scalp about 250–300 msec after feedback.

We recorded the electroencephalogram from 19 right-handed participants (9 females, aged 18–28 [M = 22.2, ±2.8]) freely navigating a “virtual” T-maze to find monetary rewards (reward: 5 cents, no-reward: 0 cents). Beginning the experiment, a robotic arm positioned the TMS coil over the left DLPFC (electrode location F3) (Fig. 1A), and continuously tracked this position to ensure precise pulse delivery (< 1 mm). Each participant received iTBS or cTBS on separate sessions (counterbalanced), with each session consisting of four blocks of 100 trials. Following the first block, 600 pulses of either iTBS or cTBS were delivered at 80% resting motor threshold. Subjects then complete three post-TBS blocks (duration 15-20 minutes).

An analysis of the reward positivity revealed a significant quadratic interaction between Block and TBS, F_1, 18_ = 6.7, p < .01, η2 = 0.21. Specifically, the delivery of iTBS following Blk1^−Baseline^ (M = −6.63 μV, ±.8) strongly reduced the reward positivity at Blk2^−postTBS^ (M = −4.80 μV, ±.7; t(18) −2.78, p<.01 ^Cohen’s d = .63^), Blk3^−postTBS^ (M = −4.66±.7 μV; t(18) −2.7, p<.05 ^Cohen’s d = .47^), and Blk4^−postTBS^ (M = −5.0 μV, ±.7; t(18) −2.2, p<.05 ^Cohen’s d = .50^). No differences were observed across cTBS blocks, and we have since replicated these results. To note, the reward positivity at baseline was comparable to other non-TBS studies (see SOM), and the amplitudes of other prominent ERP components (N2, P2 and P3) were not significantly impacted by TBS, confirming that the iTBS effect was isolated to the reward positivity.

Consistent with previous work, participants adopted a win-stay (M = 70%, +4, t(18) 5.04, p<.001) and lose-shift strategy (M = 62%, +2, t(18) −3.66, p<.005), and this strategy was maintained across blocks and did not differ between TBS protocols (Figure 2B). Further, a significant quadratic contrast was observed for reaction time for the iTBS session, F_1, 18_ = 6.5, p < .05, η2 = 0.27, with a nonsignificant linear contrast F_1, 18_ =0.73, p = .40, η2 = 0.04,. Conversely, a significant linear contrast was observed for the cTBS session, F_1, 18_ = 7.01, p < .01, η2 = 0.28, with a nonsignificant quadratic contrast F_1, 18_ = .67, p = .42, η2 = 0.04. These results suggest that reaction time constantly decreased following cTBS, but for the iTBS session, reaction time became slower and more variable across behavioral strategies following the second block. To note, aMCC lesions typically results in longer and more variable reaction times^6^, suggestive that iTBS induced a “virtual” lesion. Finally, a multiple regression analysis indicated that a decrease in reward positivity amplitude (relative to baseline) strongly predicted a decrease in selecting the alternative maze alley following feedback across Blk3^−postTBS^ and Blk4^−postTBS^ (see Figure 2C). This pattern of results was not observed in the cTBS session and no other brain – behavior relationships were observed.

**Fig 2.**
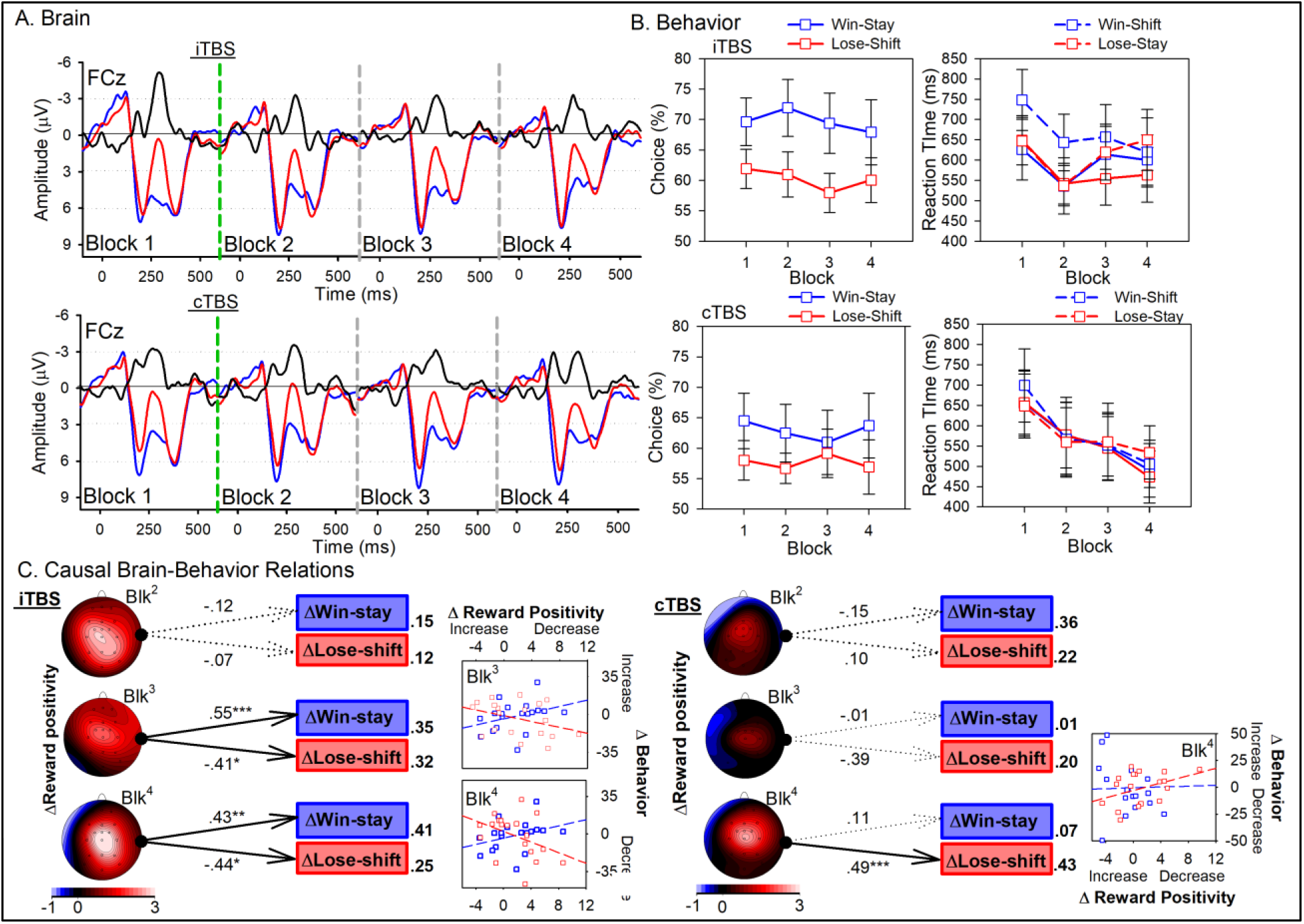
TBS results. ERP waveforms (A) and task behavior (B) associated with iTBS (top panels) and cTBS sessions (bottom panels) across blocks. ERPs elicited by reward feedback (blue), no-reward feedback (red) and difference wave (black-reward positivity). (C) Results of the multiple regression analysis for the iTBS (left) and cTBS (right) session and associated correlation plots. Predictors and DVs reflect a change in reward positivity amplitude (topoplots) and behavior (boxes) post-iTBS blocks relative to baseline. Significant direct effects and their associated beta values are shown in solid black lines. *p < 0.05, **p < 0.005, ***p < 0.001 (two-tailed). R-squared values presented right of each box.

The prominent feature of TBS over other conventional rTMS protocols is its ability to produce inhibitory (cTBS) and excitatory (iTBS) effects on motor cortex physiology using a shorter application time and lower stimulation intensity^1^. Here, we report the first evidence that reward positivity amplitude – which is said to reflect phasic modulation of aMCC activity by RPEs – was attenuated following iTBS, but not cTBS, in a simple decision-making task. Although surprising, a closer examination of the literature reveals the after-effects of TBS on prefrontal cortex function are often in conflict with motor cortex studies^4,5^. Nevertheless, such decreases in reward positivity amplitude most likely reflect either a net decrease in dopamine release to aMCC ^7^, or disrupted aMCC response to RPE signaling^9^. While the precise mechanisms of such effects are unknown, the iTBS effect on aMCC-dopamine interface is consistent with several sources of evidence. PET imaging studies indicate that left DLPFC stimulation can modulate dopamine release in aMCC ^7^; and the reward positivity, but not of other ERP components, is predictably altered by pharmaceutical and neuropsychological insults of the dopamine system ^6,10^. Alternatively, animal studies show that iTBS, but not cTBS, can induce a profound effect on cortical inhibition, which could itself cause dysfunction of cortical networks ^11^ (e.g., frontocingulate system) and possibly perturb reward processing in aMCC. Together, these interpretations may provide a common framework for relating TBS effects on aMCC-related electrophysiological signals in humans and nonhuman animals, and warrant future investigations.

Understanding the after-effects of TBS is critical if this technology is to be effectively implemented in cognitive neuroscience and psychiatric research. For instance, current theories of aMCC function — derived primarily from correlative techniques such EEG and fMRI — span a plethora of functions that include conflict processing, reinforcement learning, and error detection^12^. Because TMS offers a powerful tool for investigating causal brain-behavior relations^2^, understanding its modulatory role may help reconcile or dispute theories of aMCC function. Here, we show that decreasing reward positivity amplitude consequently reduced participants choice to select the alternative alley following feedback, regardless of its valence. One interpretation is that iTBS disrupted the aMCC valuation of the policy itself (switch alley locations following feedback), and by doing so, decreased the pursuit of such effortful behaviors. This interpretation is more in line with aMCC’s role in the valuation and selection of extended, goal-directed behavior^6^, rather than simple stimulus-response associations that characterize response-conflict and trial-and-error learning accounts of aMCC function^12^. In parallel, a substantial body of evidence implicates aMCC hypoactivity in depression patients^10^, and it has been argued that modulating cingulate activity via DLPFC might mediate the therapeutic effects of TMS^13^. If the goal of TMS in depression is to increase aMCC excitability, then applying iTBS may actually compound the impairment. However, in psychiatric conditions afflicted by aMCC hyperactivity (e.g. obsessive compulsive disorder^10^, and drug-cue hypersensitivity observed in addiction ^9^), then iTBS may prove to be beneficial.

In conclusion, instead of a general excitatory or inhibitory effect of TBS, it is reasonable to assume that TBS modulates prefrontal systems in a more complex manner than previously thought^5^. More importantly, our studies so far have now revealed both an excitatory (10 Hz rTMS)^9^ and suppressive (iTBS) action on aMCC activity, and thereby open an exciting new era of investigative possibilities in the understanding of aMCC function and treatment of aMCC dysfunction.

## Methods

### Participants

Twenty young adults (9 females, aged 18–28 [M = 22.3, SD = 2.7]) participated in the present study. All participants were right-handed (Laterality index = 70, SD = 7.5) as defined by the Edinburgh Handedness Inventory ^14^. All participants provided written informed consent and were screened for TMS permissibility, current use of Central Nervous System drugs (antidepressants, anxiolytics, antipsychotics, etc.) and recent / present neurologic symptoms using standardized forms. Participants were compensated for their participation at the rate of $15 per-hour, totaling approximately $80 USD for two experimental sessions spanning a 2 week interval. Participants were paid in full at the end of the final session and in the event that a subject wished to discontinue their participation, they were paid to the nearest half-hour. All experimental sessions took place at the Center for Molecular and Behavior Neuroscience, Rutgers-Newark. All participants had normal or corrected-to-normal vision, and were naive about the purpose of the study. The study was approved by the local research ethics committee at Rutgers University, USA, and was conducted in accordance with the ethical standards prescribed in the 1964 Declaration of Helsinki.

### Procedure

#### Virtual T-Maze task

The virtual T-maze is a reward-based choice task that elicits robust reward positivities^9,15^. In brief, at the start of each trial, participants were presented with an image of the base arm of a T-maze showing the length of the arm and two alleys projecting to the left and to the right from its far end (Figure 1C). Participants were instructed to choose one of the two arms by pressing either a left or right button. Then, they were shown a view of the alley that they selected, followed by an image of either an apple or an orange appearing against the far wall of that alley. Participants were told that presentation of one type of fruit indicated that the alley they selected contained 5 cents (reward feedback) and that the presentation of the other fruit indicated that the alley they selected was empty (no-reward feedback). Participants were informed that, at the end of the experiment, they would be rewarded all the money they found and that they were to respond in a way that maximized the total amount of money earned. Unbeknownst to them, on each trial, the type of feedback stimulus was selected at random (50% probability for each feedback type). The experiment consisted of four blocks of 100 trials each separated by rest periods (Figure 1A). Following the first block of trials, subjects received 600 pulses of TBS (see methods below). Following their last pulse (< 30 seconds), subjects continued with the three post-TBS blocks of trials.

At the end of the session, participants were asked to fill out the Alcohol, Smoking and Substance Involvement Screening Test (ASSIST),^16^ and the Substance Use Risk Profile Scale (SURPS)^17^. Participants who scored above 36 on the Global Continuum of Substance Risk (GCR) were excluded from further analysis. The data of one participant was excluded because of a history of substance misuse. Following the questionnaires, subjects were debriefed about the purpose of the study and were compensated for their participation.

#### T-maze Behavioral Analysis

The percentage of choice (e.g. win-stay and lose-shift) and the reaction time were calculated for behavioral analyses. All trials with reaction time faster than 100ms or slower than 3000ms were excluded from the analyses. Specifically, if subjects chose the same or different alley on the trial N after receiving reward (i.e. apple) on the previous trial, the trial N is defined as a Win-Stay or Win-Shift trial. Similarly, if subjects chose the same or different alley on the trial N after receiving no-reward (i.e. orange) on the previous trial, the trial N is defined as a Lose-Stay or Lose-Shift trial. The percentage of Win-Stay or Win-Lose choice were calculated based on the number of Win-Stay or Win-Lose trials divided by the total of Win-Stay and Win-Shift trials. Similarly, the percentage of Lose-Shift choice were calculated based on the number of Lose-Shift trials divided by the total of Lose-Shift and Lose-Stay trials. The averaged reaction time on these four trial types (i.e. Win-Stay, Win-Shift, Lose-Stay, and Lose-Shift) were also calculated.

#### EEG Data Acquisition and Analysis

Continuous EEG was recorded from 27 scalp electrodes (Fp1, Fp2, F7, F3, Fz, F4, F8, FC5, FC1, FCz, FC2, FC6, T7, C3, Cz, C4, T8, CP5, CPz, CP6, P3, Pz, P4, PO7, POz, PO8, Oz), placed according to the 10/20 system. Signals were acquired using Ag–AgCl ring electrodes mounted in a nylon electrode cap with a conductive gel (Falk Minow Services, Herrsching, Germany). Signals were amplified by low-noise electrode differential amplifiers with a frequency response of DC 0.017–67.5 Hz (90-dB octave roll-off) and digitized at a rate of 1000 samples per second. Digitized signals were recorded to disk using Brain Vision Recorder software (Brain Products GmbH, Munich, Germany). Electrode impedances were maintained below 10 kΩ. Two electrodes were also placed on the left and right mastoids. The EEG was recorded using the average reference.

#### ERP Analysis

Postprocessing and data visualization were performed using Brain Vision Analyzer software (Brain Products GmbH). The digitized signals were filtered using a fourth-order digital Butterworth filter with a bandpass of .10–20 Hz. An 1-sec epoch of data extending from 200 msec before to 800 msec after the onset of each feedback stimulus was extracted from the continuous data file for analysis. Ocular artifacts were corrected using the eye movement correction algorithm described by Gratton, Coles, and Donchin (1983). The EEG data were rereferenced to linked mastoids electrodes. The data were baseline-corrected by subtracting from each sample the mean voltage associated with that electrode during the 200-msec interval preceding stimulus onset. Muscular and other artifacts were removed using a ±150-μV level threshold and a ±35-μV step threshold as rejection criteria. ERPs were then created for each electrode and participant by averaging the single-trial EEG according to feedback type (Reward, No-reward) and Block (Block 1 [pre-TBS baseline], Block 2 [post-TBS 1], Block 3 [post-TBS 2], Block 4 [post-TBS 3]).

##### Reward Positivity

Reward positivity amplitude is typically assessed as the size of the difference in the ERPs elicited to positive and negative feedback stimuli of equal expectancy ^9,15^. Thus, to isolate the reward positivity from other overlapping ERP components, the reward positivity was evaluated for each electrode channel and participant as a difference wave by subtracting the Reward feedback ERPs from the corresponding No-reward feedback ERPs. As before^9^, reward positivity amplitude was determined at FCz by identifying the maximum absolute amplitude of the difference wave within a 200-to 400-ms window following feedback onset for block (Blk1^−Baseline^, Blk2^−postTBS^, Blk3^−postTBS^, Blk4^−postTBS^).

##### Other ERP components

A consideration of TBS modulators’ effects is also valuable in interpreting additional ERP components in the post-feedback waveform, particularly the P200 (Table S1), N200 (Table S2), and P300 (Table S3). The P200 was measured using a local peak detection method by identifying the maximum positive value of the ERP within a window extending from 100 ms to 250 ms following the presentation of the feedback, the time of which was taken as the peak latency of the P200. This algorithm was applied to the Reward and No-reward ERPs associated with electrode site FCz. The N200 was quantified at FCz and its amplitude was measured for each participant and feedback condition (reward, no reward) using a local peak detection algorithm, in which the most negative value within a time window starting from 200 to 400 ms after feedback presentation was taken as the peak. P300 amplitude was measured by identifying the maximum positive-going value of the Reward and No-reward ERPs recorded at electrode site Pz, within a window extending from 300 to 600 ms following the presentation of the feedback stimulus.

#### Statistical analysis strategy

##### Reward Positivity

For the purpose of statistical analysis, reward positivity was evaluated at channel FCz where it reaches is maximal. To test our predictions, we conducted a two-factor mixed-design analysis of variance (ANOVA) on reward positivity amplitude with block (Blk1^−Baseline^, Blk2^−postTBS^, Blk3^−postTBS^, Blk4^−postTBS^) and TBS session (iTBS, cTBS) as within-subject factors. Sex, age, and TBS order were controlled for throughout our analysis. The Greenhouse– Geisser correction for nonsphericity was applied where appropriate, and the Benjamini– Hochberg procedure was used to correct for multiple testing in all post hoc analyses.

##### Behavior

For reaction time, we conducted a four-factor mixed-design ANOVA on reaction time with block (Blk1^−Baseline^, Blk2^−postTBS^, Blk3^−postTBS^, Blk4^−postTBS^), feedback (win, lose), behavior (stay, switch), and TBS session (iTBS, cTBS) as within-subject factors. For percentage of choice, we conducted a three-factor mixed-design ANOVA with block (Blk1^−Baseline^, Blk2^−postTBS^, Blk3^−postTBS^, Blk4^−postTBS^), strategy (win-stay, loose-shift) and TBS session (iTBS, cTBS) as within-subject factors. Sex, age, and TBS order were controlled for throughout our analysis. The Greenhouse–Geisser correction for nonsphericity was applied where appropriate, and the Benjamini–Hochberg procedure was used to correct for multiple testing in all post hoc analyses.

##### Brain and Behavior relationships

A multiple regression analysis was performed using the computer program AMOS 18.0.1 (Arbuckle, 1995–2009) to identify unique relationships between a change in reward positivity amplitude and a change in behavior following iTBS and cTBS ([Blk1^−Baseline^ -Blk2^−postTBS^], [Blk1^−Baseline^ - Blk3^−postTBS^], [Blk1^−Baseline^ - Blk4^−postTBS^]). Type 1 errors were statistically controlled following Benjamini and Hochberg (1995) with a corrected significance level of α = 0.05. Sex, age, and TBS order were included in the regression model as nuisance variables.

#### Robot-assisted TMS protocol

Robotic-assisted TMS is a novel approach in the application for TBS protocols. For each subject, a 3D model of the head surface was created in Smartmove (ANT, Enschede, Netherlands). A headmarker carrying three passive Polaris reflective markers on a plastic frame was fixed to the head and tracked by a Polaris infrared stereo-optical tracking system. The construction of the surface of the 3D head model was done by pointer acquisition of up to 600 surface points. This also determined the coordinate transformation from the robot end-effector to the coil. The robot and the tracking system were registered to a common coordinate system. To set the target for TBS, the left DLPFC target (here, electrode location F3), the coil was placed on the surface of the 3D head model, and the corresponding point on the surface of the virtual head model was selected and the orientation of the coil was calculated tangentially to cortical surface and 45° to a sagittal plane. The distance between the TMS coil and the selected DLPFC target was less than 10 mm. The virtual coil position was then transformed to robot coordinates, which defined the movement of the robot to the corresponding target position relative to the subject’s head. The position of the cranium was continuously tracked and the trajectory of the coil’s path was adapted to movements of the head by a motion compensation module in real-time, which guaranteed a precise coil to target position and allowed free head movements during the study^18^.

#### TBS protocol

The stimulator device was a MagPro X100 with the Cool-B70 figure-of-eight coil (MagVenture, Falun, Denmark). Resting Motor Threshold (rMT) measurements were performed via visual twitch in the contralateral (right) hand. The coil was positioned over the supposed motor cortex area, then the coil was moved until the location at which a reproducible abductor pollicis brevis response, elicited at the lowest stimulator intensity could be identified. rMT was defined as the lowest stimulation intensity, expressed as a percentage of max output of the Magstim equipment that reliably (3/5 times) yielded a visible muscle twitch in the hand when stimulating the hand area of the contralateral motor cortex with a single pulse, and that was set as rMT. In keeping with safety guidelines, stimulation intensity for the experiment was set for each individual participant as 80% of motor threshold (range 55% - 75%). Average stimulation intensity was 53% (range: 44%– 56%) of maximum output.

Participants received iTBS or cTBS over the predefined left DLPFC target (F3) on separate occasions, with order counterbalanced across participants. For each session, participants completed 400 trials of the virtual T-maze task. Following the first block of 100 trials, participants received 600 pulses of either iTBS or cTBS. Following Huang et al. (2005) protocol, cTBS involved an uninterrupted train of TBS for 40s, in which the inhibitory after-effects can last for up to 50min post-stimulation. Conversely, iTBS was comprised a 2s TBS train (10 TBS bursts) delivered every 10s (8s inter-burst interval between trains) over a total of 190s, resulting in an excitatory effect for up to 60min post stimulation. As noted above, stimulation intensity for each TBS session was set for each participant as 80% of motor threshold as evaluated on the first session. One person could not tolerate the TBS protocol, and the experiment was stopped. In total, 20 participants completed both TBS sessions. Following TBS, participant then completed 3 blocks of 100 trials, with each block lasting approximately 5-7 minutes. After the task was completed, the EEG cap was removed and participants were debriefed and compensated for their participation in the study.

## Supplementary material

**Figure S1.**
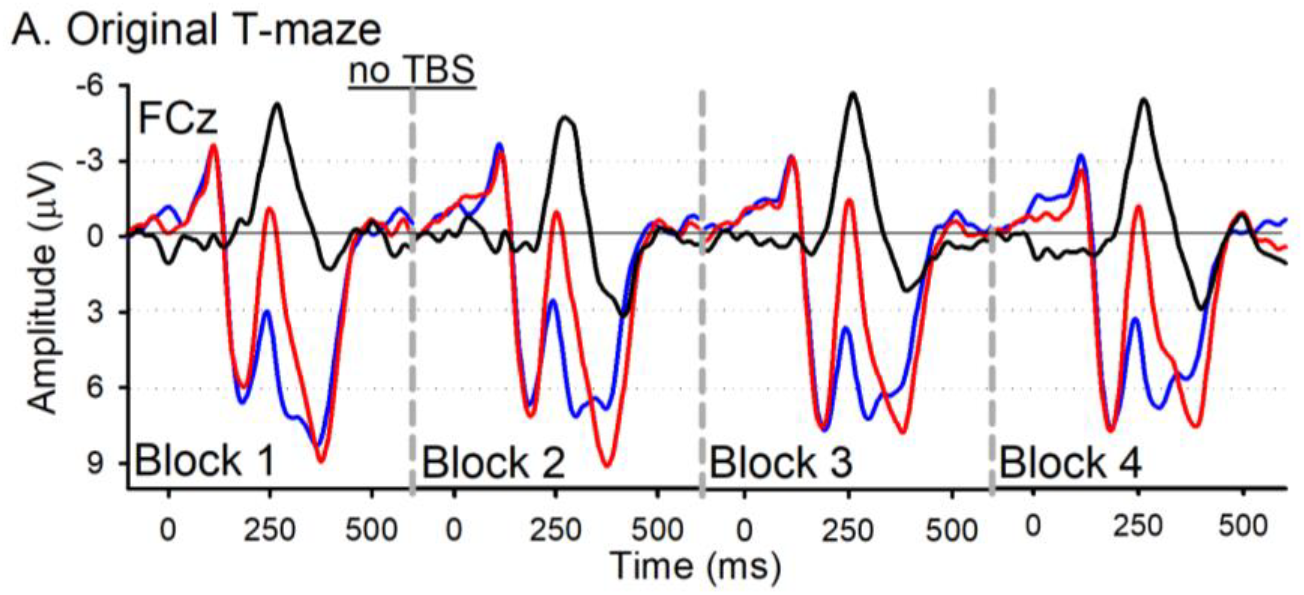
ERP waveforms from a previous study collected without TBS (data adapted from Baker et al., 2009). ERPs elicited by reward feedback (blue), no-reward feedback (red) and difference wave (black-reward positivity) across 4 blocks of trials (100 trials per block). To note, the reward positivity did not differ between Block 1 and the subsequent blocks (p>.05).

**Table S1.** Mean and standard deviation of P2 peak amplitude for cTBS and iTBS under Reward and No-reward conditions for each block (window: 160-260ms post-feedback)

|  | Block 1 | Block 2 |  | Block 3 |  | Block 4 |  |
| --- | --- | --- | --- | --- | --- | --- | --- |
|  | *M*<br>(*SD*) | *M*<br>(*SD*) | *p* value<br>(vs.<br>block1) | *M*<br>(*SD*) | *p* value<br>(vs.<br>block1) | *M*<br>(*SD*) | *p* value<br>(vs.<br>block1) |
| **No-reward** |  |  |  |  |  |  |  |
| cTBS (µV) | 6.63<br>(2.82) | 7.76<br>(2.74) | 0.04 | 7.54<br>(2.84) | 0.07 | 7.93<br>(3.56) | 0.03 |
| iTBS (µV) | 8.51<br>(2.44) | 9.14<br>(3.02) | 0.17 | 8.89<br>(3.82) | 0.57 | 8.32<br>(3.61) | 0.77 |
| *p* value<br>(cTBS vs. iTBS) | 0.01 | 0.06 |  | 0.04 |  | 0.67 |  |
| **Reward** |  |  |  |  |  |  |  |
| cTBS (µV) | 8.59<br>(4.08) | 8.88<br>(3.42) | 0.56 | 9.06<br>(4.21) | 0.42 | 9.36<br>(4.29) | 0.18 |
| iTBS (µV) | 8.58<br>(2.54) | 9.61<br>(3.19) | 0.01 | 9.23<br>(3.90) | 0.39 | 8.73<br>(3.92) | 0.83 |
| *p* value<br>(cTBS vs. iTBS) | 0.98 | 0.30 |  | 0.81 |  | 0.35 |  |

**Table S2.** Mean and standard deviation of N2 peak amplitude for cTBS and iTBS under Reward and No-reward conditions for each block (window: 200 - 400ms post-feedback).

|  | **Block 1** | **Block 2** |  | **Block 3** |  | **Block 4** |  |
| --- | --- | --- | --- | --- | --- | --- | --- |
|  | *M*<br>(*SD*) | *M*<br>(*SD*) | *p* value<br>(vs.<br>block1) | *M*<br>(*SD*) | *p* value<br>(vs.<br>block1) | *M*<br>(*SD*) | *p* value<br>(vs.<br>block1) |
| **No-reward** |  |  |  |  |  |  |  |
| cTBS (μV) | -1.62<br>(3.04) | -1.87<br>(4.06) | 0.75 | -1.87<br>(3.53) | 0.72 | -0.65<br>(3.12) | 0.15 |
| iTBS (μV) | -1.73<br>(3.92) | -0.76<br>(3.85) | 0.09 | -1.20<br>(3.75) | 0.38 | -1.04<br>(3.69) | 0.22 |
| *p* value<br>(cTBS vs. iTBS) | 0.90 | 0.24 |  | 0.39 |  | 0.62 |  |
| **Reward** |  |  |  |  |  |  |  |
| cTBS (μV) | 1.32<br>(3.83) | 1.31<br>(3.48) | 0.99 | 0.82<br>(4.25) | 0.30 | 1.36<br>(3.93) | 0.95 |
| iTBS (μV) | 2.33<br>(2.94) | 1.44<br>(3.43) | 0.11 | 1.50<br>(3.47) | 0.25 | 1.46<br>(3.39) | 0.26 |
| *p* value<br>(cTBS vs. iTBS) | 0.35 | 0.90 |  | 0.35 |  | 0.90 |  |

**Table S3.**
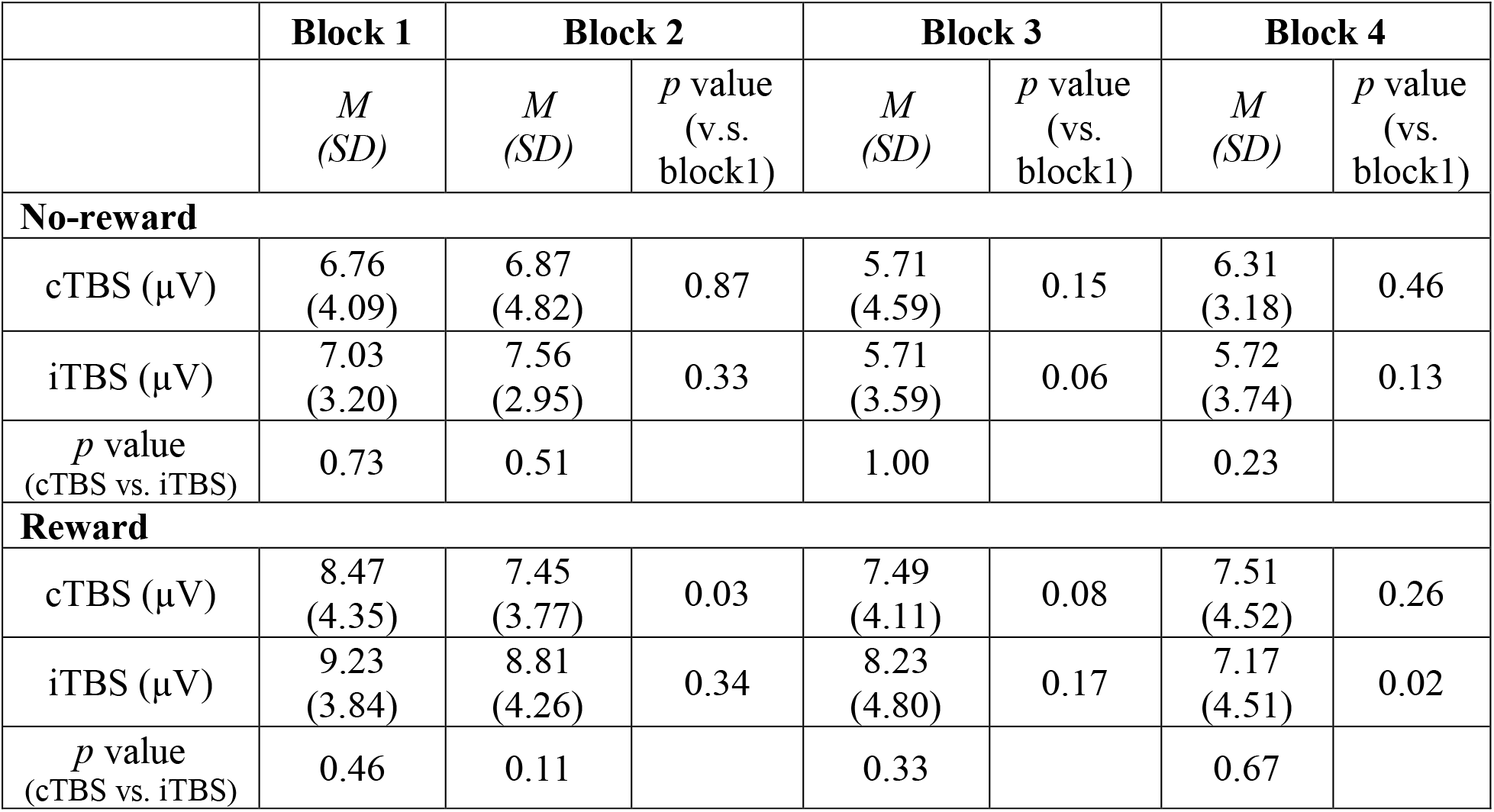
Mean and standard deviation of P3 peak amplitude for cTBS and iTBS under Reward and No-reward conditions for each block (window: 300 – 460ms post-feedback)

|  | **Block 1** | **Block 2** |  | **Block 3** |  | **Block 4** |  |
| --- | --- | --- | --- | --- | --- | --- | --- |
|  | *M*<br>(*SD*) | *M*<br>(*SD*) | *p* value<br>(v.s.<br>block1) | *M*<br>(*SD*) | *p* value<br>(vs.<br>block1) | *M*<br>(*SD*) | *p* value<br>(vs.<br>block1) |
| **No-reward** |  |  |  |  |  |  |  |
| cTBS (μV) | 6.76<br>(4.09) | 6.87<br>(4.82) | 0.87 | 5.71<br>(4.59) | 0.15 | 6.31<br>(3.18) | 0.46 |
| iTBS (μV) | 7.03<br>(3.20) | 7.56<br>(2.95) | 0.33 | 5.71<br>(3.59) | 0.06 | 5.72<br>(3.74) | 0.13 |
| *p* value<br>(cTBS vs. iTBS) | 0.73 | 0.51 |  | 1.00 |  | 0.23 |  |
| **Reward** |  |  |  |  |  |  |  |
| cTBS (μV) | 8.47<br>(4.35) | 7.45<br>(3.77) | 0.03 | 7.49<br>(4.11) | 0.08 | 7.51<br>(4.52) | 0.26 |
| iTBS (μV) | 9.23<br>(3.84) | 8.81<br>(4.26) | 0.34 | 8.23<br>(4.80) | 0.17 | 7.17<br>(4.51) | 0.02 |
| *p* value<br>(cTBS vs. iTBS) | 0.46 | 0.11 |  | 0.33 |  | 0.67 |  |

**Table S4.** Mean and standard deviation of Win-Stay and Lose-Shift choice (%) for cTBS and iTBS by block

|  | **Block 1** | **Block 2** |  | **Block 3** |  | **Block 4** |  |
| --- | --- | --- | --- | --- | --- | --- | --- |
|  | *M*<br>(*SD*) | *M*<br>(*SD*) | *p* value<br>(vs.<br>block1) | *M*<br>(*SD*) | *p* value<br>(vs.<br>block1) | *M*<br>(*SD*) | *p* value<br>(vs.<br>block1) |
| **Win-Stay Choice** |  |  |  |  |  |  |  |
| cTBS (%) | 64<br>(20) | 62<br>(21) | 0.53 | 61<br>(23) | 0.46 | 64<br>(23) | 0.91 |
| iTBS (%) | 70<br>(17) | 72<br>(20) | 0.39 | 69<br>(22) | 0.93 | 68<br>(23) | 0.61 |
| *p* value<br>(cTBS vs. iTBS) | 0.27 | 0.05 |  | 0.09 |  | 0.42 |  |
| **Lose-Shift Choice** |  |  |  |  |  |  |  |
| cTBS (%) | 58<br>(14) | 57<br>(11) | 0.53 | 59<br>(17) | 0.71 | 57<br>(19) | 0.76 |
| iTBS (%) | 62<br>(14) | 61<br>(16) | 0.73 | 58<br>(14) | 0.29 | 60<br>(16) | 0.69 |
| *p* value<br>(cTBS vs. iTBS) | 0.37 | 0.33 |  | 0.69 |  | 0.44 |  |

**Table S5.** Mean and standard deviation of reaction time on Win-Stay, Win-Shift, Lose-Stay, and Lose-Shift trials for cTBS and iTBS by block

|  | **Block 1** | **Block 2** |  | **Block 3** |  | **Block 4** |  |
| --- | --- | --- | --- | --- | --- | --- | --- |
|  | *M*<br>(*SD*) | *M*<br>(*SD*) | *p* value<br>(vs.<br>block1) | *M*<br>(*SD*) | *p* value<br>(vs.<br>block1) | *M*<br>(*SD*) | *p* value<br>(vs.<br>block1) |
| **Win-Stay** |  |  |  |  |  |  |  |
| cTBS (ms) | 655.69<br>(355.35) | 575.14<br>(411.14) | 0.11 | 549.96<br>(362.08) | 0.08 | 490.39<br>(288.12) | 0.04 |
| iTBS (ms) | 626.00<br>(326.05) | 536.11<br>(302.24) | 0.01 | 614.08<br>(298.87) | 0.82 | 600.13<br>(271.36) | 0.61 |
| *p* value<br>(cTBS vs. iTBS) | 0.71 | 0.58 |  | 0.28 |  | 0.06 |  |
| **Win-Shift** |  |  |  |  |  |  |  |
| cTBS (ms) | 699.26<br>(382.84) | 566.86<br>(384.32) | 0.01 | 552.27<br>(326.05) | 0.02 | 505.90<br>(246.31) | 0.02 |
| iTBS (ms) | 747.58<br>(327.52) | 643.50<br>(296.01) | 0.10 | 656.73<br>(338.57) | 0.21 | 619.22<br>(361.09) | 0.18 |
| *p* value<br>(cTBS v.s. iTBS) | 0.52 | 0.45 |  | 0.05 |  | 0.21 |  |
| **Lose-Stay** |  |  |  |  |  |  |  |
| cTBS (ms) | 648.82<br>(342.80) | 559.02<br>(373.21) | 0.21 | 560.43<br>(415.26) | 0.30 | 533.82<br>(286.26) | 0.09 |
| iTBS (ms) | 646.63<br>(253.78) | 537.10<br>(216.98) | 0.07 | 619.71<br>(300.55) | 0.64 | 649.86<br>(327.06) | 0.96 |
| *p* value<br>(cTBS vs. iTBS) | 0.98 | 0.78 |  | 0.42 |  | 0.07 |  |
| **Lose-Shift** |  |  |  |  |  |  |  |
| cTBS (ms) | 656.62<br>(339.68) | 577.09<br>(354.57) | 0.11 | 544.72<br>(339.90) | 0.09 | 473.40<br>(279.27) | 0.02 |
| iTBS (ms) | 648.52<br>(264.76) | 541.98<br>(218.28) | 0.04 | 554.30<br>(285.94) | 0.10 | 563.60<br>(294.32) | 0.14 |
| *p* value<br>(cTBS vs. iTBS) | 0.91 | 0.64 |  | 0.84 |  | 0.14 |  |

